# Plant cellulose synthase membrane protein isolation directly from *Pichia pastoris* protoplasts, liposome reconstitution, and its enzymatic characterization

**DOI:** 10.1101/2023.03.29.534738

**Authors:** Dharanidaran Jayachandran, Shoili Banerjee, Shishir P. S. Chundawat

## Abstract

The most abundant renewable biopolymer on earth, viz., cellulose, acts as carbon storage reserve in plant and microbial cell walls that could potentially be converted into biofuels or other valuable bioproducts. Cellulose is synthesized by a plant cell membrane-integrated processive glycosyltransferase (GT) called cellulose synthase (CesA). Since only a few of these plant CesAs have been purified and characterized to date, there are huge gaps in our mechanistic understanding of these enzymes. Furthermore, the coordination between different CesAs involved in primary and secondary cell wall formation is yet to be unveiled. The biochemistry and structural biology studies of CesAs are currently hampered by challenges associated with their expression and extraction at high yields. To aid in understanding CesA reaction mechanisms and to provide a more efficient CesA extraction method, two putative plant CesAs – PpCesA5 from *Physcomitrella patens* and PttCesA8 from *Populus tremula x tremuloides* that are involved in primary and secondary cell wall formation in plants were expressed using *Pichia pastoris* as an expression host. We developed a protoplast-based membrane protein extraction approach to directly isolate both these membrane-bound enzymes for purification, as detected by immunoblotting and mass spectrometry-based analyses. Our method results in a higher purified protein yield by 3-4-fold than the standard cell homogenization protocol. Our purified CesAs were reconstituted into liposomes to yield active enzymes that gave similar biochemical characteristics (e.g., substrate utilization and cofactor requirements, no primer needed to initiate polymerization reaction) as enzymes isolated using the standard protocol. This method resulted in reconstituted CesA5 and CesA8 with similar Michaelis-Menten kinetic constants, K_m_ = 167 μM, 108 μM and V_max_ = 7.88×10^−5^ μmol/min, 4.31×10^−5^ μmol/min, respectively, in concurrence with the previous studies. Taken together, these results suggest that CesAs involved in primary and secondary cell wall formation can be expressed and purified using a simple and more efficient extraction method. This could potentially help unravel the mechanism of native and engineered cellulose synthase complexes involved in plant cell wall biosynthesis.

## 1. Introduction

Polysaccharides are a major class of natural polymers found in the plant, animal, and microbial kingdoms that are essential in providing energy, structural support, and other biological functions [1–3]. These complex carbohydrates are synthesized by a group of enzymes called polysaccharide synthases [4–7]. Some of these polysaccharide synthases are membrane-integrated processive family-2 glycosyltransferases, such as cellulose, hyaluronan, chitin, and alginate synthases [8]. Polysaccharides such as cellulose and hemicellulose are the most abundant renewable polymers found in plant cell walls. Cellulose, an unbranched homopolysaccharide made up of D-glucose linked by β-1,4-glycosidic bonds [9], is the major structural component of plant cell walls and is also found in algae and some microbes. It is used in several industries, including but not limited to paper, textiles, and furniture. In recent decades, cellulose and its associated proteins have gained much attention since it could be used as a potential feedstock for producing bioethanol and other valuable bioproducts [10–13]. Therefore, increasing plants’ biomass yield and sugar content is imperative and possible by altering the cell wall composition [14]. However, understanding the fundamental mechanisms and factors influencing the formation of these polysaccharides is far from fruition.

Cellulose is processively synthesized by a membrane-integrated processive family-2 glycosyltransferase called cellulose synthase (CesA). These enzymes exist in nature as membrane-localized complexes [15–17] and typically contain multiple monomers that coordinate amongst themselves and carry out various biological functions. For instance, in *Arabidopsis thaliana*, CesAs interact to form rosette subunits, and six of these subunits assemble into multimeric rosette complexes, often called cellulose synthase complexes (CSCs). These CSCs contain several different CesA isoforms that express differentially during various stages of cell wall formation [18,19]. Arabidopsis expresses ten different *CesA* genes with different subsets that are involved in either primary cell wall formation (proteins encoded by *CesA*1, *CesA*3, and *CesA*6 or *CesA*2/5/9) or secondary cell wall formation (proteins encoded by *CesA*4, *CesA*7, and *CesA*8) [20]. Recently, the structure of a homotrimeric CSC containing three CesA8 monomers from Poplar was solved using CryoEM, which revealed a molecular basis for understanding cellulose microfibril formation [17]. Each CesA monomer comprises seven transmembrane helices circumscribed by the intracellular N- and extracellular C-terminus and a large cytosolic GT domain. Likewise, the homotrimeric structure of CesA7 from cotton was also resolved in a similar manner and showed an analogous structure [21].

Plasma membrane-localized CSCs are made up of different individual CesA isoforms that are responsible for processively synthesizing single glucan chains and assembling them into the cellulose microfibril (CMF) matrix. Recent biochemical studies show that a single CesA isoform, when functionally reconstituted into a liposome, is enough to synthesize cellulose microfibrils or form UDP as a bi-product when incubated with UDP-glucose as substrate [22–24].

Although seminal research in the last couple of years has revealed the structure of plant CesA and its activity *in vitro*, there are still significant gaps in our understanding of how these CesA monomers coordinate together and form microfibrils both *in vivo* and *in vitro*. It is important to note that such an imperative plant protein system has only a few reports available on their expression and purification to date. This is mainly due to the lack of reports that elucidate simple, efficient, and feasible methods of expression and purification. In this work, we intend to showcase an efficient method of purification that could potentially help prepare and study different CesAs side by side. To achieve this, we selected two putative CesAs (CesA5 from *Physcomitrella patens* and CesA8 from *Populus tremula x tremuloides*) involved in the primary and secondary cell wall formation. Both these enzymes were expressed heterologously in *Pichia pastoris* and purified using a modified protoplast extraction method, as confirmed by various detection methods. The enzymes were reconstituted into proteoliposomes and produced UDP when incubated with UDP-Glucose as a substrate and manganese as a cofactor. We have performed steady-state kinetic analysis and determined various kinetic parameters for both the enzymes. To our knowledge, this is the first study in which enzymes involved in both primary and secondary plant cell wall formation have been studied side by side *in vitro*. Overall, our results show that the modified extraction approach is suitable for both CesA5 and CesA8 without impacting the catalytic activity.

## 2. Materials and methods

### 2.1 Cloning and transformation into yeast

Cellulose synthase 8 (*CesA*8) gene from hybrid aspen (*Populus tremula x tremuloides*) carrying a C-terminal dodeca-HIS-tag and an N-terminal FLAG tag [17], and Cellulose synthase 5 (*CesA*5) gene from moss (*Physcomitrella patens*) carrying a C-terminal dodeca-HIS-tag [23] was custom synthesized from GenScript and cloned into yeast expression vector pPICZA. Plasmid maps for both constructs are shown in Figure S1. Protein sequences for CesA5 and CesA8 are shown in Supplementary text S1. The construct was then transformed into the *Pichia pastoris* SMD1168H strain (single protease deficient strain) using the Easyselect Pichia Expression kit (Invitrogen, Cat# K174001) according to the manufacturer’s specifications. The cells were plated on YPDS plates [1% yeast extract, 2% peptone, 2% dextrose, 1 M sorbitol, 2% agar (w/v)] containing 100 μg/mL zeocin and were incubated for 2-4 days at 30 °C. The colonies were screened using colony PCR by checking the integration of the gene into alcohol oxidase (AOX I) loci using primers specific to the AOX promoter (5’ AOX 1 primer: 5’-GACTGGTTCCAATTGACAAGC-3’ and 3’ AOX 1 primer -5’-GCAAATGGCATTCTGACATCC-3’). All the primers used in this study were obtained from Integrated DNA Technologies and are tabulated in Supplementary Table S1.

### 2.2 Growth conditions for expression of CesA proteins

Growth conditions were similar to the one mentioned in Purushotham et al., with slight modifications [22]. Transformed cells were cultured overnight at 30°C in 5 ml YPDS culture tubes. Approximately 3-5% of this culture was inoculated into 300 ml preculture media (BMGY medium containing 100 mM Phosphate buffer pH 6.0, 1% yeast extract, 2% peptone, 1.34% yeast nitrogen base, 1% glycerol) and incubated overnight at 30°C and 300 rpm. Cells were collected after 12-16 hours and resuspended to an OD_600_ of 0.4 in BMMY induction media (BMGY medium supplemented with 0.5% methanol instead of glycerol). Induction was carried out in baffled flasks at 20°C and 300 rpm for 24 h. The cells were later harvested at 7000 rpm for 20 mins, and the cell pellets were directly used for purification or stored at −80°C for long-term storage.

### 2.3 Extraction and purification of enzymes using Pichia protoplasts

Traditional methods of protein extraction from yeast, such as sonication, bead-beating, and homogenization, were employed to move the membrane protein (MP) from the cell surface to the soluble fraction. However, the sonication or the bead beating methods were futile since the both of them resulted in no yield at all. For homogenization method, we resuspended 12g of harvested cells from 1L culture in 60 mL lysis buffer (20 mM Tris-HCl pH 7.5, 0.6 M sorbitol) and lysed by two passes through a homogenizer at ∼15,000 psi in the presence of one cOmplete™ EDTA-free protease inhibitor tablet per 50 mL sample volume. The lysate was centrifuged at 19,000 × g for 10 min at 4 °C, and the supernatant was centrifuged for 2 h at 100,000 × g and 4 °C to pellet the membrane fraction. The membrane pellet was solubilized in 60 mL membrane resuspension buffer (MRB) (20 mM Tris-HCl pH 7.5, 100 mM NaCl, 40 mM n-Dodecyl β-D -Maltoside (DDM), 10% vol/vol glycerol, and one cOmplete™ EDTA-free protease inhibitor tablet) and incubated at 4 °C for 120 mins with gentle agitation. Insoluble material was removed by centrifugation at 100,000 × g for 30 min at 4 °C. Supernatant was then subjected to IMAC based purification as mentioned in the next section. However, the high-pressure homogenizer method resulted in a very low yield of CesA proteins (Fig. S2).

Hence, we used a gentler method involving a short solubilization step directly on whole cells to favor the extraction of the undamaged, correctly folded MPs targeted to the plasma membrane [25]. In this method, 12g of harvested cells from 1L culture was initially washed with 200 mL double-distilled water to remove any residual media, followed by 200 mL SED buffer (1M Sorbitol, 25 mM EDTA, and 1M DTT). The cells were later washed using 200 mL of 1M sorbitol before resuspending them in 150 mL of CG buffer (20 mM trisodium citrate pH 5.8, 10% glycerol, 1 mM PMSF). 20 units of zymolyase (from Amsbio, UK) per gram of cells was added to the mixture and incubated at 30°C and 70 rpm for 20-30 mins. The resulting yeast spheroplasts or protoplasts (yeast cells without a cell wall) were later used to directly solubilize membrane proteins in 100 mL solubilization buffer (containing 50 mM Tris–HCl pH 7.4, 500 mM NaCl, 10% glycerol, 20 mM imidazole, 40 mM DDM, and cOmplete™ EDTA-free protease inhibitor tablet). After solubilizing the membrane proteins for 2-2.5 h at 4°C with gentle agitation, the samples were centrifuged at 48,400 x g at 4°C for 1 h in a Beckman Coulter JA-20 fixed angle rotor. The supernatant was collected and filtered using a 0.22 µm non-sterile syringe filter and incubated with 5 ml preequilibrated TALON superflow resin (Cytiva, Cat# 28957502) overnight at 4°C with gentle agitation. The schematic workflow of the traditional and protoplast-based lysis methods is shown in Fig. 1.

**Figure 1.**
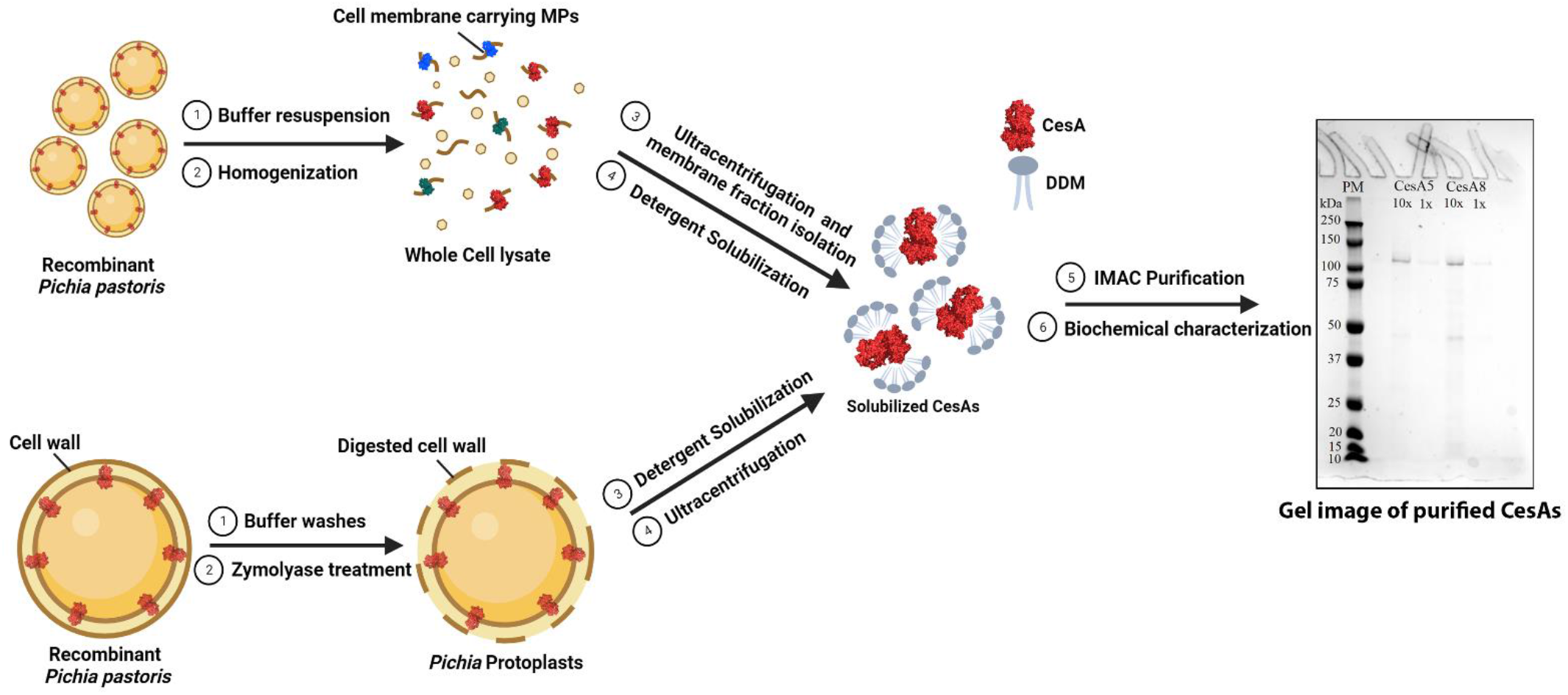
Schematic workflow of CesA purification methods. (*Top*) Recombinant Pichia cells expressing the CesA (shown in red) lysed using the homogenization method followed by membrane fraction extraction, detergent solubilization using N-dodecyl β-D-maltoside (DDM), and purification. (*Bottom*) Protoplasts-based extract method through multiple buffer washes (double-distilled water, SED, Sorbitol) and Zymolyase treatment. Zymolyase digests cell walls, forming protoplasts that were then directly used for protein solubilization using DDM. The gel on the right is a Coomassie-stained SDS PAGE gel depicting 1x and 10x concentrated CesA5 and CesA8. Proteins depicted in red are CesA8 (PDB:6WLB). MP: Membrane proteins. The image is not to scale.

### 2.4 Gravity-based IMAC purification of CesA5 and CesA8

The resin was packed into a gravity flow column and sequentially washed with equilibration (EQ) buffer (20 mM Tris–HCl pH 7.5, 100 mM NaCl, 10% glycerol) containing 20-, 40-, or 60-mM imidazole and 1 mM LysoFoscholineEther-14. For CesA5, an additional washing step with EQ buffer containing 80 mM imidazole and 1 mM LysoFoscholineEther-14 was required before the final elution. CesA proteins were eluted in the EQ buffer containing 300 mM imidazole. The eluted fraction was concentrated 10 times using 100 kDa Amicon Ultra-15 centrifugal filters (Millipore Sigma, Cat# UFC903008) and buffer exchanged onto a pre-equilibrated PD-10 column (Cytiva, Cat# 17085101). The concentration of the purified protein was estimated using BCA assay (ThermoFisher, Cat# 23225). Samples were stored at 4°C and reconstituted immediately. For long-term storage, it is recommended to flash freeze the samples and store them at -80°C.

### 2.5 Immunoblot analysis

After subjecting the samples to polyacrylamide gel electrophoresis (SDS-PAGE), the proteins were carefully transferred onto a nitrocellulose membrane (Bio-Rad, Cat# 1620112) of dimensions (8.6 × 6.7 cm) at 100V with a constant current for 60 mins at 4°C in a Bio-Rad Mini-Transfer Cell (Bio-Rad, Cat# 1703930) according to the manufacturer’s specifications. The nitrocellulose membrane was blocked with 3% (w/v) BSA/PBS-Tween 20 solution overnight at 4°C. After blocking for 14-16 hours, the membrane was washed six times for 5 mins with gentle agitation at 25°C with PBS/Tween 20 buffer. The membrane was then incubated for 1 h with anti–His primary mouse antibodies (1:1000) at room temperature. The membrane was then washed thrice for 5 mins in PBS-Tween 20 before incubation with an HRP-conjugated anti-mouse secondary antibody (1:1000) and streptactin antibody (specific to the standard protein marker – Bio-Rad, Cat# 1610376; 1:1000 dilution) for 1 h at room temperature. After washing the membrane six more times, the membrane was incubated with clarity western ECL substrate (Bio-Rad, Cat#1705060) and imaged using a chemiluminescence imager (Syngene Pxi 4 EZ).

### 2.6 Reconstitution of cellulose synthase into liposomes

4 mg/ml of yeast total lipid extract (Avanti polar lipids, Cat# 190000C) was taken using glass Pasteur pipets in clean glass vials (Avanti polar lipids, Cat# 600460), and the chloroform was entirely removed by blowing it with a stream of nitrogen gas. The vials were kept under vacuum overnight in a desiccator to remove any residual organic solvent. The lipid was later solubilized in 400 µl EQ buffer containing 120 mM LDAO by vortexing vigorously and placing it in running warm water until the solution got clear. 600 µl of concentrated protein was added to this mixture and incubated on ice for an hour to form mixed micelles. Meanwhile, 5g of SM2 bio-beads (Bio-Rad, Cat# 1523920) were washed for 5 mins, twice with 50 ml methanol and thrice with 50 ml water using a magnetic stirrer before storing them at 4°C in DI water. These washed bio-beads were dried at room temperature on tissue paper/Kim wipes for 10 mins before using. Dried bio-beads were added sequentially to the reconstitution mixture to prevent aggregate formation. 0.35 g of bio-beads was added to the reconstitution mixture and incubated at 4°C for 1 h with gentle agitation. After an hour, the sample was transferred to a fresh vial containing 0.35 g of bio-beads, and the mixture was incubated overnight at 4°C with gentle agitation. On the next day, bio-beads were allowed to settle under gravity, and the supernatant was pipetted out carefully without disturbing the beads. The supernatant was then subjected to ultracentrifugation at 60,000 rpm (∼200,000xg) in a Beckman Coulter fixed angle rotor (TLA 100.3) for 45 mins at 4°C. The supernatant was discarded, and the pellet containing liposomes was washed with 1 ml EQ buffer (without detergent). The suspended liposomes were subjected to ultracentrifugation (∼200,000xg for 45 mins at 4°C) to dilute any residual detergent. The supernatant was then discarded, and the liposome pellet was resuspended in 1 ml EQ buffer (without detergent). This sample was subjected to extrusion using Avanti mini extruder (Avanti polar lipids, Cat# 6100001EA) fitted with a 100 nm pore size filter to form uniformly sized vesicles. The extrusion was performed 15-21 times before collecting the samples. The extruded sample was subjected to a final ultracentrifugation step at ∼200,000xg for 30 mins at 4°C to remove aggregates. The supernatant was then collected, and the samples were stored at 4°C before carrying out the activity assay. For long-term storage, samples were aliquoted and flash-frozen before storing them at -80°C.

### 2.7 Cellulose synthase activity assays

Standard cellulose synthase assays were set up according to Omadjela et al [26]. Twenty microliters of PttCesA8-or PpCesA5 containing proteoliposomes were incubated in the presence of 10 mM MnCl_2_, 3 mM UDP-Glucose, in a buffer containing 20 mM Tris (pH 7.5), 100 mM NaCl, and 10% (vol/vol) glycerol. After incubation at 37°C for 3 h, the samples were centrifuged at 15,000 rpm for 20 mins. 10 µl supernatant was incubated with 10 µl of freshly prepared nucleotide detection reagent for UDP-Glo assays (Promega, Cat#: V6961) according to manufacturer’s specifications. The samples were incubated at room temperature for an hour, and luminescence was recorded using luminescence protocol in a Spectramax M5 plate reader. All the studies were performed in triplicates, and the error bars reported are standard deviation from the mean.

### 2.8 Time course and kinetic studies of reconstituted cellulose synthases

To analyze the time taken for the CesAs to reach saturation, the reconstituted proteoliposomes were incubated with 3 mM UDP-Glucose, and 20 mM MnCl_2_ for 4 h at 37°C and samples were collected at regular intervals before running the UDP-Glo assay as mentioned previously. Alternatively, for characterizing the kinetics of CesAs, the samples were incubated in the presence of 20 mM MnCl_2_ and 0–3.5 mM UDP-Glucose. A stock concentration of 300 mM UDP-Glucose was used to dilute the substrate concentration in each reaction vial. After synthesis for 30 mins, the reaction mixture was subjected to UDP-Glo assay as described above. All the studies were performed in triplicates, and the error bars reported are standard deviation from the mean.

### 2.9 Kinetic analysis and calculations

Preliminary data analysis was performed using Microsoft Excel™ to obtain UDP produced (μmol/min). The data was fit to the monophasic Michaelis-Menten kinetic tool in Origin to obtain V_max_ and K_m_. The turnover number (k_cat_) was calculated from V_max_ using the following equation, as outlined in detail elsewhere [27]:

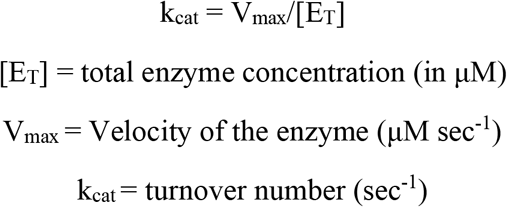

Similarly, the data obtained from the time-course study was fitted using the non-linear curve fitting tool in Origin. Curve fitting was done using the Levenberg-Marquardt algorithm with a tolerance of 1e-9.

## 3. Results and Discussion

### 3.1 Heterologous expression and cell lysis for the extraction of CesAs

Cellulose synthase was predicted to have seven transmembrane helices, an N-terminal Zn-binding domain, a large cytosolic domain with a TED motif, and plant-conserved and class-specific regions [22,23]. When the structure of the homotrimeric CesA8 from Poplar was resolved using Cryo-EM, these predictions became more transparent [17]. Here, we used the same set of genes reported previously, but the codon-optimized versions of PpCesA5 conjugated with C-terminal 12x His-tag and PttCesA8 conjugated with N-terminal FLAG tag, and C-terminal 12x HIS-tag for heterologous expression in *Pichia pastoris*. CesA genes were integrated into the genome of Pichia under the control of the AOX1 promoter. Hence, the induction of the protein was performed using methanol as an inducer. The integration of the gene and its whole sequence was confirmed using the primers listed in Supplementary Table S1.

Membrane proteins (MP) are highly amphipathic, temperature, and shear-sensitive. Conventional methods of membrane protein extraction, such as bead-beating, sonication, product entrapment, and homogenization, exert a lot of physical pressure on the cells that could be detrimental to the membrane protein integrity [28–30]. Moreover, methods like homogenization might also extract the MPs that are folded incorrectly and not processed completely since it involves the usage of whole-cell lysate [25]. During this study, the high-pressure homogenizer or the bead beating method was fruitless since the former resulted in a very low yield of purified CesA proteins (Fig. S2), and the latter resulted in no yield. The yield of CesAs from homogenized protein samples was found to be between 25-40 µg/ml from a 1L batch. Hence, we used a slightly modified Pichia protoplast-based extraction method that is gentler on the cells and involves chemical and enzymatic treatment followed by a short solubilization step directly on protoplasts to favor the extraction of the undamaged, correctly folded MPs that have been targeted to the plasma membrane (Fig. 1). This could potentially help overcome the problem of decreased yields in CesAs since the probability of getting correctly folded MPs is more. We observed a 3-4-fold increase in the amounts of both CesA5 and CesA8 when we used the modified protoplast extraction approach compared to homogenization. Only a tiny fraction of the proteins recovered from the total lysate were CesAs (Supplementary Table S2).

### 3.2 Purification and detection of PttCesA8 and PpCesA5

CesAs were purified to homogeneity in the detergent Lysofoscholine Ether 14 (LFCE14) via immobilized metal affinity chromatography (IMAC). The isolated membrane fraction obtained from the modified protoplast extraction method was directly used for purification. The different fractions involved in the purification of PpCesA5 and PttCesA8 were observed under protein detection techniques such as Coomassie and silver staining. The enriched proteins were found to be immunoreactive when treated with anti-HIS antibodies (Fig. 2; Fig. S3). The final elute had highly enriched PttCesA8 and PpCesA5 proteins of approximately 110 and 125 kDa, respectively, when compared to a standard protein marker. A ∼50 kDa band was observed under SDS-PAGE analysis in both cases. Interestingly, this band did not appear when the fraction was raised against the anti-HIS antibody, as mentioned in some previous reports [22,23]. Also, reducing the zymolyase treatment time from 30 mins to 20 mins nearly removed the ∼50 kDa band observed in the case of both CesAs (Fig. S4). Longer exposure to zymolyase treatment could have resulted in delicate protoplasts making it more susceptible to cell lysis and protein degradation. Relative quantity and percentage purity were determined by analysis of SDS–PAGE band intensities using the Image Lab software, version 6.0.1 (Bio-Rad) as mentioned elsewhere [31]. The percentage purity was 75.8 and 79.2 for both CesA5 and CesA8. Also, the relative quantity of CesAs in the purified elute compared to membrane solubilized fraction was 55.35 and 52.85 respectively. These values for both CesA5 and CesA8 are tabulated in Supplementary Table S3.

**Figure 2.**
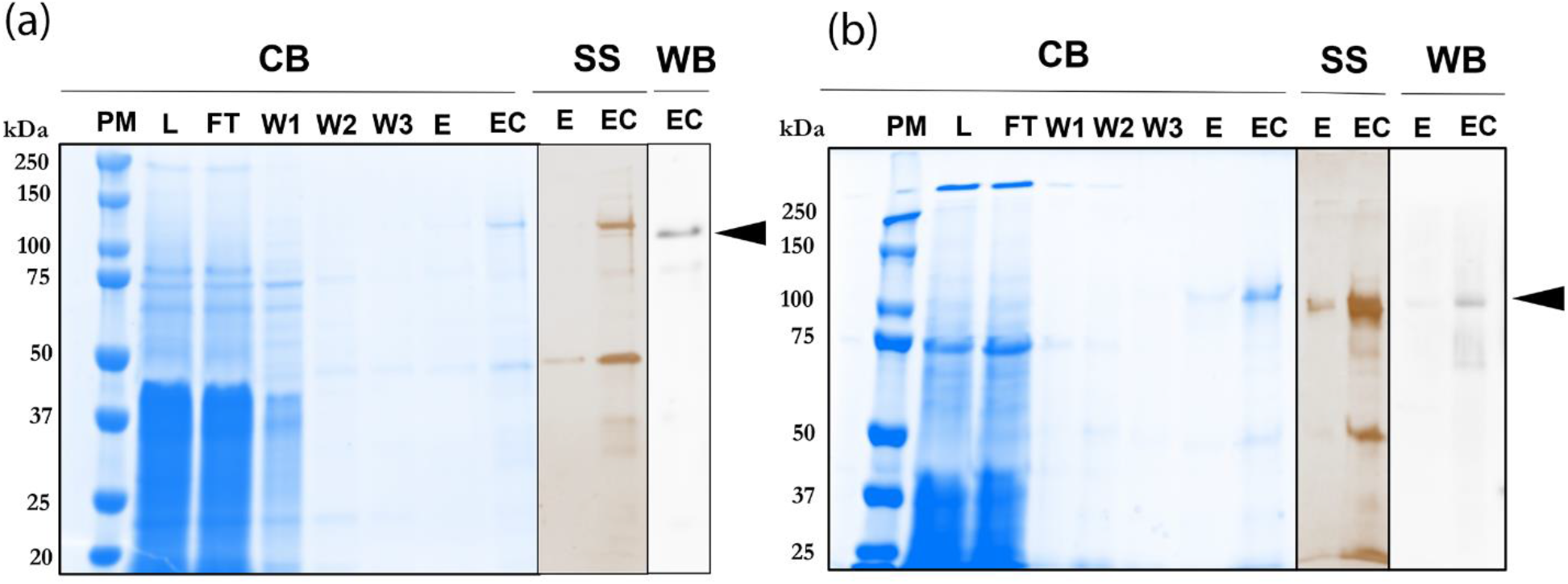
Coomassie blue (CB)- and Silver (SS)-stained SDS-PAGE WB –Western Blot-raised against the C-terminal His-tag of (a) PpCesA5 (b) PttCesA8. PM-Protein Marker; L-Load; FT-flow-through; W1–3, wash steps 1–3; E, eluted fraction; EC, 10x Concentrated eluted fraction. The black arrowheads represent the position of the purified CesA enzymes.

Although Coomassie, silver staining, and immunoblotting detected the presence of HIS-tagged CesAs, we wanted to confirm the presence of PpCesA5 and PttCesA8 further using LC-MS-MS. The bands corresponding to the molecular weight of both PpCesA5 and PttCesA8 were excised and analyzed at a tandem mass spectrometry fingerprinting facility at Rutgers. Thirty-seven peptides specific to PpCesA5 and fifty-four peptides specific to PttCesA8 were identified, confirming the presence of both these proteins (Supplementary Tables S4 and S5). The other proteins observed using mass spectrometry were mostly contaminating proteins arising from the expressing organism. None of those contaminating proteins shows any documented evidence or function in polysaccharide biosynthesis. Interestingly, no peptides matching *Pichia* β-1,3 glucan synthase were observed using this method, contrary to the homogenizer based methods reported previously [22,23].

### 3.3 Time course study shows a faster saturation for CesA5 compared to CesA8

Purified CesAs were reconstituted into yeast total lipid extract liposomes using the detergent-mediated liposome reconstitution method [22]. The reconstituted enzyme’s catalytic activity was measured in the presence of 3 mM UDP-Glucose and 20 mM Mn^2+^. Reactions catalyzed by CesAs result in the formation of UDP nucleotide that could be measured to quantify CesA activity. Control reactions in the absence of proteoliposomes did not contain any UDP and the background was subtracted from the obtained values. As shown in Fig. 3a and b, PpCesA5, and PttCesA8 continued to produce UDP at an optimal pH of 7.5 and temperature of 37°C. The catalytic activity stalls after 180 min of incubation for PttCesA8 and after 120 min of incubation for PpCesA5, respectively. This could be due to the depletion in protein activity or inhibition of catalytic activity by UDP produced. The maximum amount of UDP produced was 84 nM in the case of PpCesA5 and 72 nM in the case of PttCesA8. A similar trend in saturation was observed for the previously reported time-course synthesis studies of CesAs [22–24]. The time course study for both the enzymes were observed to fit into a non-linear model as shown in supplementary Fig. S5. The fitted curve shows the maximum UDP produced (P1) and the time taken to produce half of the maximum UDP (P2). The curve is observed to follow a non-linear trend before flattening out completely indicating saturation of product accumulation. Value of P2 in CesA5 is roughly double the time as that of CesA8 suggesting a faster accumulation of UDP in CesA5 compared to CesA8.

**Figure 3.**
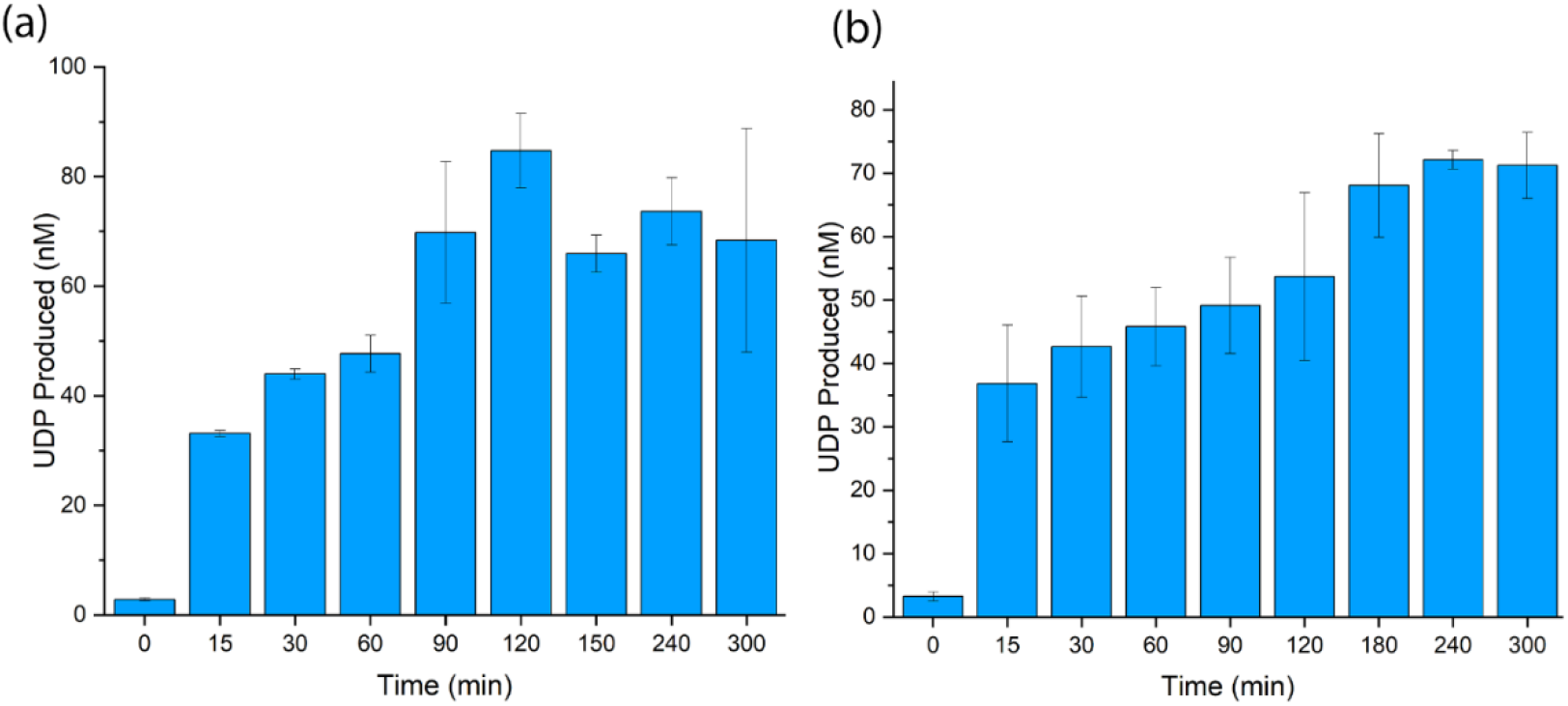
Time course of UDP biosynthesis using UDP-glucose as a substrate by reconstituted (a) PpCesA5 (b) PttCesA8.

**Figure 4.**
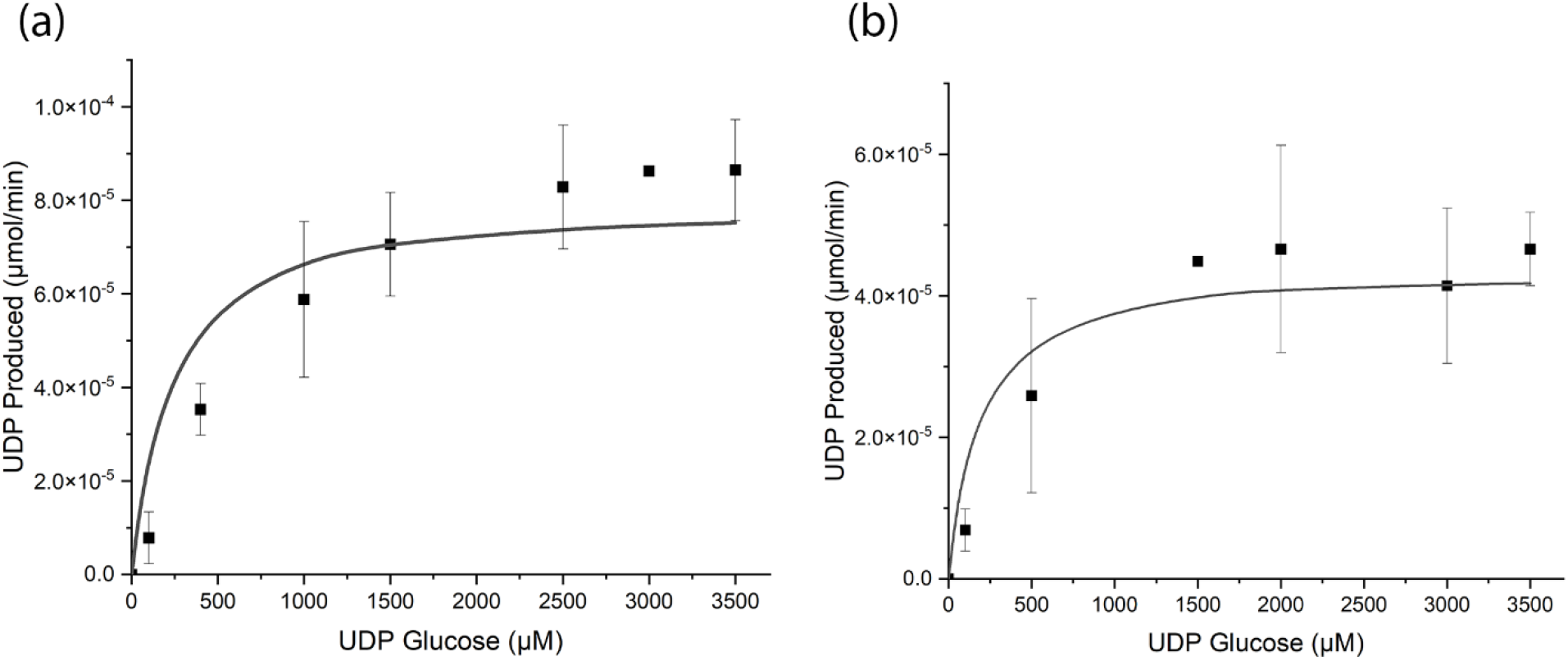
Kinetic analyses of (a) PpCesA5 and (b) PttCesA8 by titrating increasing amounts of UDP-Glucose and quantification of UDP. The obtained data were fit to monophasic Michaelis-Menten kinetics using Origin software, yielding a *K*_*m*_ of 167 µM and 108 µM.

The time-course activity data was eventually used to calculate the specific activity of the proteoliposomes (Supplementary table S2). The specific activity of membrane solubilized fraction was observed to be the highest (161.62 and 157.97 nmol/min/mg for CesA5 and CesA8) since the UDP produced could be from other contributing fungal enzymes like β-1,3 glucan synthase that utilize UDP-Glucose as substrate [32–34]. Interestingly, these enzymes were not observed in neither of our purification preparations when observed under mass spectrometry (Supplementary Tables S4 and S5). Hence, the UDP formed from Co-TALON elute fraction and reconstituted liposomes is mostly from CesAs. The higher value observed from the reconstituted liposomes could be attributed to the greater stability the lipid vesicles provide to the membrane proteins than detergent micelles.

Compared to the bacterial cellulose synthases, the plant cellulose synthase shows almost a two-magnitude difference in the specific activity according to one report [35] and similar specific activity in another [36]. However, the values reported previously were obtained directly from the purified and total membrane fractions and not from the functionally reconstituted liposomes. It would therefore be premature on our part to make a direct comparison between the specific activities across two different types of samples from two different species.

### 3.4 Reconstituted CesA8 shows a higher substrate affinity compared to CesA5

To measure the affinity of the substrate towards the enzymes, we measured the apparent K_m_ values for both the reconstituted proteoliposomes. K_m_ values of PpCesA5 and PttCesA8 were estimated to be 167 µM, and 108 µM, consistent with the values reported for reconstituted CesAs previously [22–24]. Also, a comparable K_m_ value of 500 μM is observed in the case of reconstituted *R. sphaeroides* BcsA [26] and 270 μM in the case of AcsA-B from *G. hansenii* [37]. All the data were fit to monophasic Michaelis–Menten kinetics, and a lower K_m_ value in the case of CesA8 suggests a greater affinity of the substrate towards this enzyme compared to CesA5.

The turnover number (k_cat_) of CesA5 and CesA8 was calculated to be 1.45 ± 0.13 sec^-1^ and 0.79 ± 0.06 sec^-1^ respectively. Correspondingly, the catalytic efficiency was estimated to be 0.009 ± 0.01 µM^-1^ sec^-1^ and 0.007 ± 0.01 µM^-1^ sec^-1^ for each of the enzymes. All the kinetic parameters are summarized in Table 1. The reproducibility of the values was tested with two separate purified enzyme preparations.

**Table 1.** Summary of kinetic parameters for both CesA5 and CesA8 enzymes. Experiments were run in duplicates, and errors are standard deviations from the mean.

| Enzyme | $K_m$ ($\mu\text{M}$) | $V_{\text{max}}$ ($\mu\text{mol/min}$) | $k_{\text{cat}}$ ($\text{sec}^{-1}$) | $k_{\text{cat}}/K_m$ ($\mu\text{M}^{-1} \text{ sec}^{-1}$) |
| --- | --- | --- | --- | --- |
| CesA5 | $166.95 \pm 13.47$ | 7.88E-05 | $1.45 \pm 0.13$ | $0.009 \pm 0.01$ |
| CesA8 | $108.57 \pm 9.78$ | 4.31E-05 | $0.79 \pm 0.06$ | $0.007 \pm 0.01$ |

The k_cat_ for cellulose synthase has been published only for bacterial cellulose synthases until now. Cellulose synthase from *R. sphaeroides* has been reported to have a k_cat_ of 90 sec^-1^. Also, the *Gluconacetobacter hansenii* enzyme has been reported to have a k_cat_ of 1.60 ± 0.50 sec^-1^ which was almost two orders of magnitude lower than that of *R. sphaeroides* [26]. Interestingly, plant cellulose synthases also had k_cat_ in a similar range as that of *G. hansenii* [35,37,38], meaning they also have lower values than that of *R. sphaeroides*. It is difficult to assess whether the plant CesAs are slower in adding the substrate than *R. sphaeroides* since a recent study involving cellulose biosynthesis at a single-molecule level shows an addition of glucose every 2.5 secs at room temperature [39]. Such state-of-the-art methods may be required to determine the exact catalytic efficiency and turnover rate of the plant CesAs.

## 4. Conclusion

Primary cell walls are synthesized during cell expansion and are highly extensible and incorporative. On the other hand, secondary cell walls are not extensible and typically provide tensile strength and rigidity after the cell ceases expansion [40]. Primary CesAs are known to physically interact both *in vitro* and *in planta*, with all secondary CesAs suggesting specialized functions for CesAs in certain developmental or environmental conditions [41,42]. These CesAs typically interact and form cellulose synthase complex (CSC) in higher plants [43–45]. Therefore, a systematic study at an enzymatic level is imperative to compare and contrast the different CesAs involved in primary and secondary cell wall synthesis, respectively. In this study, we have developed a simple and efficient method of CesA extraction from recombinant Pichia protoplasts.

In conclusion, our work confirms that different CesAs involved in the primary and secondary cell wall formation extracted using the Pichia protoplast-based method are catalytically active and show similar biochemical and kinetic characteristics to some of the previous studies. This method also results in a higher purified enzymatic yield than the homogenization-based method. We also observed that some of the kinetic characteristics of plant CesAs are similar to those of bacterial CesAs. The developed method also allows access to purified, membrane-bound, functional CesAs that may yield structures of CesAs in the future. Such studies may eventually unravel the coordination between various CesAs inside CSC in vascular and non-vascular plants.

## Author Contributions

The original manuscript draft was written by DJ and edited by SPSC. DJ and SB conducted all the experiments. DJ and SPSC designed the study. All authors have given approval to the final version of the manuscript.

## Acknowledgement and Funding Sources

This work was supported by the U.S. Department of Energy (Award Number: DE-SC0019313), Rutgers Energy Institute (REI), and Rutgers University. We thank Prof. James Evans and Dr. Amar Parvate for discussions and valuable inputs. We also thank Prof. Jean Baum and Prof. Andy Nieuwkoop for help with access to the homogenizer and ultracentrifuge used in this study. We are also grateful to Prof. Jay Sy for access to the chemiluminescence imager.

## Abbreviations

CSC: Cellulose Synthase Complex;
CesA: Cellulose Synthase;
Ptt: *Populous tremuloides x tremuloides*;
Pp: *Physcomitrella patens*;
UDP: Uridine Di-Phosphate;
MP: Membrane Protein.

