## Supplementary information for "Plant cellulose synthase membrane protein isolation directly from *Pichia pastoris* protoplasts, liposome reconstitution, and its enzymatic characterization"

**This SI file includes the following:**

Figures S1 to S5 (Page 2-4)

Tables S1 to S5 (Pages 5-10)

Supplementary text S1 (Page 11)

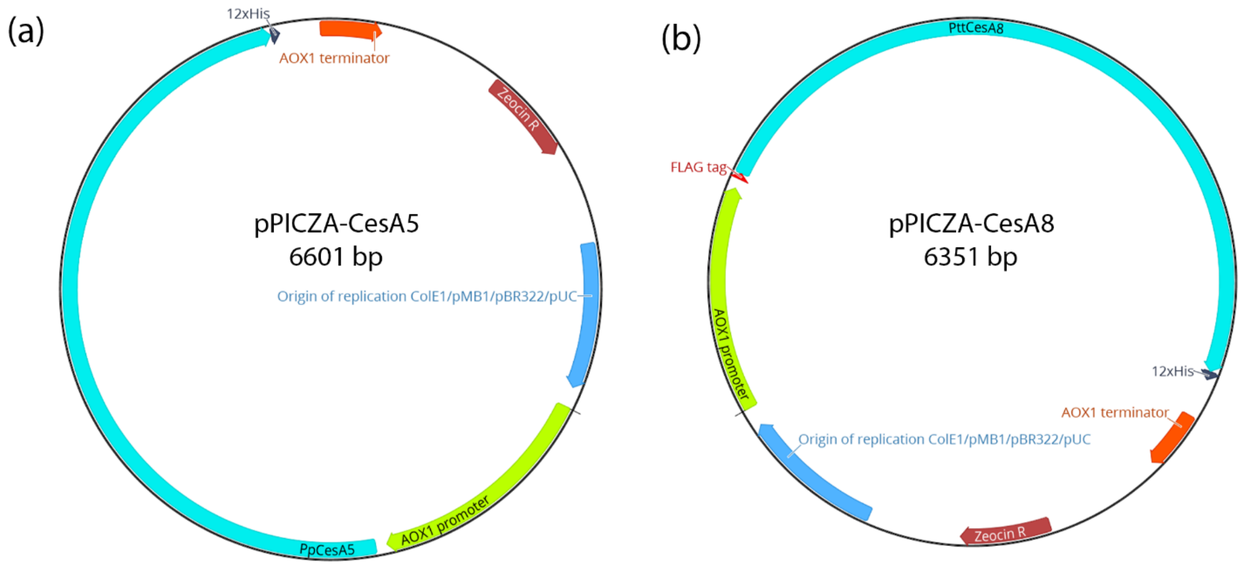


**Figure S1:** Plasmid maps for pPICZa-CesA5 and pPICZa-CesA8 indicating the AOX1 promoter region, CesA followed by 12x-HIS tag, and ending with AOX1 terminator. pPICZa-CesA8 has an additional FLAG tag after the promoter region and before the CesA8 gene.

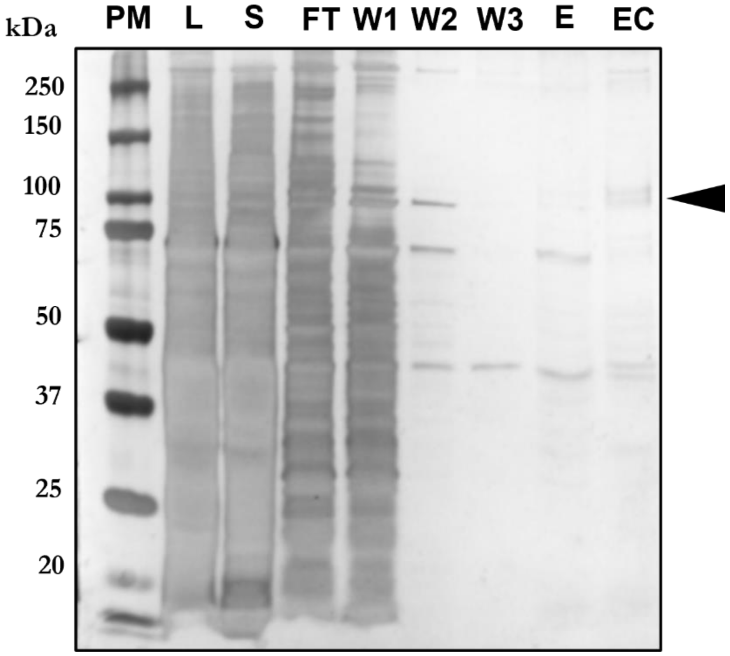


**Figure S2:** Silver stained SDS-PAGE gel of purified PttCesA8 showing various fractions of a homogenized sample. PM- Protein Marker; L- Whole cell lysate; S – Supernatant; FT- flow-through; W1–3, wash steps 1–3; E, eluted fraction; EC, 10x Concentrated eluted fraction. PpCesA5 showed no band from two different sets of experiments.

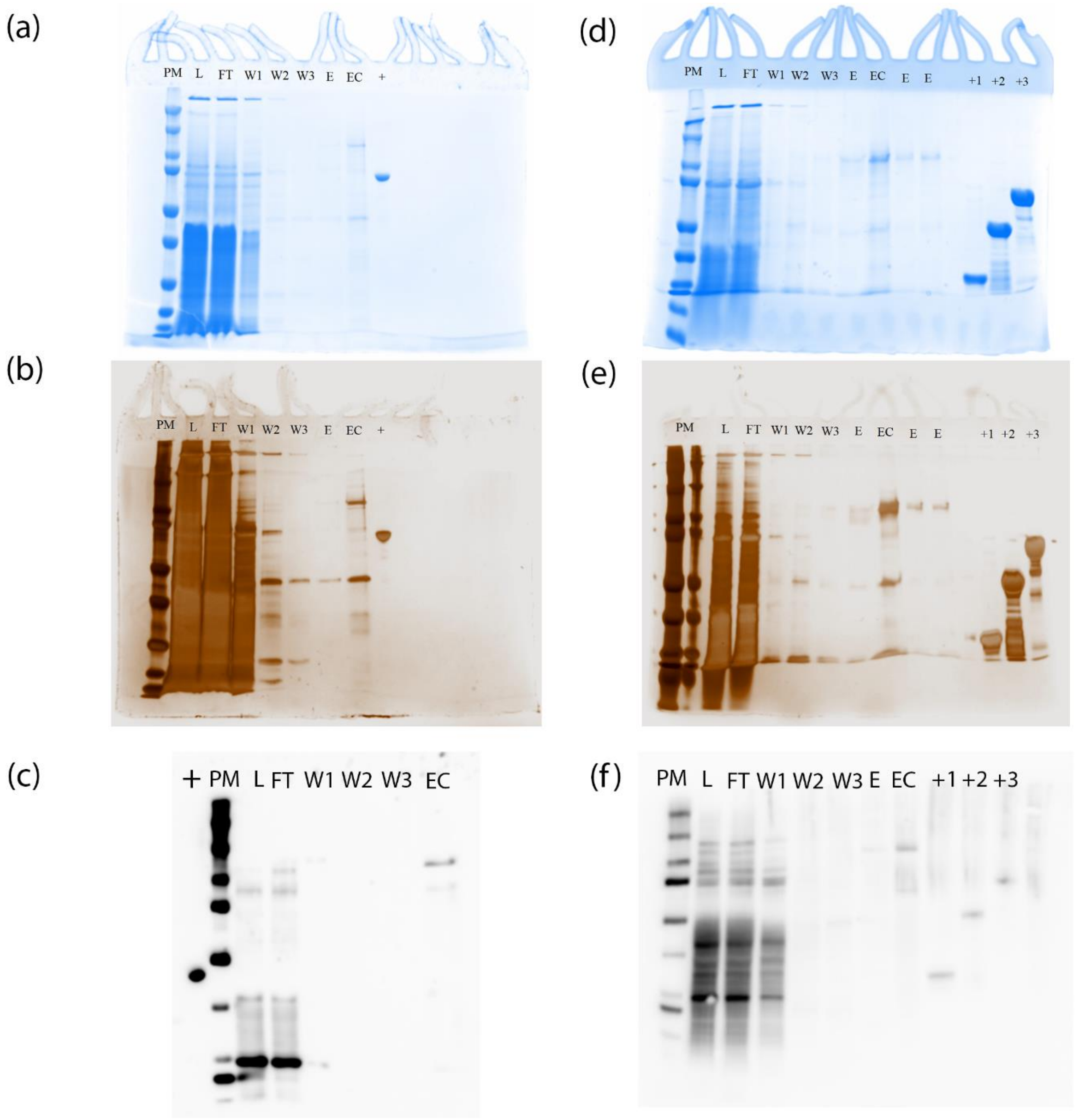


**Figure S3:** Blots and gels shown in Figure 2 without cropping. Coomassie blue (CB)- and Silver (SS)-stained SDS-PAGE WB –Western Blot- raised against the C-terminal His-tag of (a-c) PpCesA5 (d-f) PttCesA8. (+) His-tagged positive controls of different sizes made in-house.

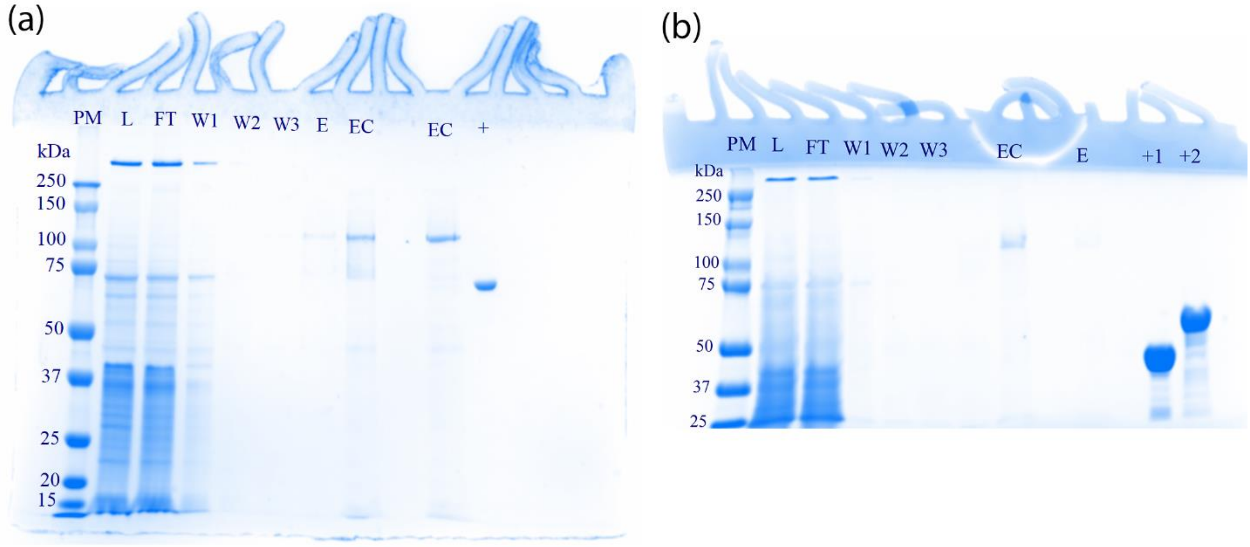


**Figure S4:** Coomassie blue (CB)- SDS-PAGE gels (a) PpCesA5 (b) PttCesA8 showing purified samples from reduced zymolyase treatment (20 mins). These gels were later used to calculate percentage purity and relative quantity. (+) His-tagged positive controls of different sizes made in-house.

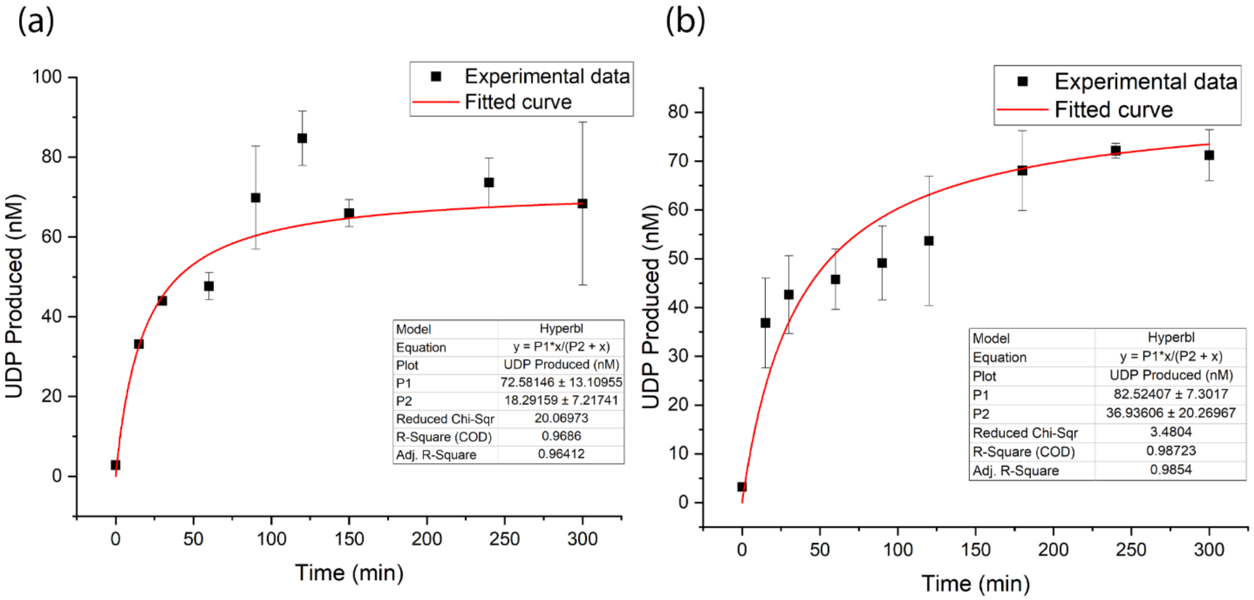


**Figure S5:** Non-linear fitted curve of CesA5 and CesA8 time-course study. The curves show saturation time points and the maximum amount of UDP produced.

**Supplementary Table S1**. Primer sequences for target ORF amplification and sequencing.

| **Name of Primer** | **Sequence** |
|---|---|
| 5’ AOX 1 primer | GACTGGTTCCAATTGACAAGC |
| 3’ AOX 1 primer | GCAAATGGCATTCTGACATCC |
| PpCesA5_For1 | CGACAACTTGAGAAGATC |
| PpCesA5_For2 | TTTGCATCAAACTCCTCAA |
| PpCesA5_For3 | AGTGGTTGCCAATTAACA |
| PpCesA5_For4 | CCTGGTATGATTCAAGTTTT |
| PpCesA5_For5 | GACTGCTCCTACTAGATC |
| PpCesA5_For6 | TCCAACTATCTCTAATTTGGA |
| PttCesA8_For1 | ACTAATTATTCGAAACGAGG |
| PttCesA8_For2 | AAACTGAACCAGCTCAAG |
| PttCesA8_For3 | AGCTGAGTTTGCAAGGAA |
| PttCesA8_For4 | ATCCACAAGTAGGTCGAG |
| PttCesA8_For5 | CTGTGGCTTTGAAGAGAA |
| PttCesA8_For6 | TTTCAGCCCATCTCTTTG |

**Supplementary Table S2.** Specific activities for CesA5 and CesA8[a]

| **Step** | | **Volume (ml)** | **Protein concentration (mg/ml)** | **Total protein in 20 µl reaction (mg)** | **Total activity (nmol/min)** | **Specific activity (nmol/min/mg)** |
|---|---|---|---|---|---|---|
| Resuspended protoplasts[b] | | 110 | - | - | - | - |
| Membrane solubilized 2h[c] | CesA5 | 100 | 3.39 | 0.068 | 10.99 | 161.62 |
| | CesA8 | 100 | 3.18 | 0.064 | 10.11 | 157.97 |
| Co-TALON IMAC | CesA5 | 3 | 0.11 | 0.002 | 0.58 | 29 |
| | CesA8 | 3 | 0.12 | 0.002 | 0.51 | 25.5 |
| Liposomes[d] | CesA5 | 1 | 0.05 | 0.001 | 0.47 | 47 |
| | CesA8 | 1 | 0.05 | 0.001 | 0.4 | 40 |

[a]Cellulose synthase activity was measured by UDP-Glo assay as mentioned under Section 2.
[b]Zymolyase-treated protoplasts resuspended in the solubilization buffer.
[c]After solubilization of MPs from protoplasts into detergent, the total membrane fraction along with protoplasts was centrifuged as described in Section 2.
[d]After reconstitution and extrusion of CesAs into liposomes, the concentration was measured with BCA assay compared to blank liposomes.

**Supplementary Table S3.** Relative quantity and purity percentage were determined by analysis of SDS–PAGE band intensities shown in Figure S4 using the Image Lab software, version 6.0.1 (Bio-Rad).

| **Samples** | | **Relative quantity** | **Percentage purity** |
|---|---|---|---|
| Membrane solubilized | CesA5 | 1 | - |
| | CesA8 | 1 | - |
| Co-TALON IMAC | CesA5 | 55.35 | 75.8 |
| | CesA8 | 52.85 | 79.2 |

**Supplementary Table S4.** List of *Pichia pastoris* proteins in the excised single band (~125 kDa) corresponding to PpCesA5 as identified by mass spectrometry.

| Protein | Description | Unique | Total |
|---|---|---|---|
| tr\|A0A1B2JF91\|A0A1B2JF91_PICPA | BA75_03713T0 {ECO:0000313\|EMBL:ANZ76707.1}; | 447 | 928 |
| tr\|A0A1B2JHN5\|A0A1B2JHN5_PICPA | BA75_05137T0 {ECO:0000313\|EMBL:ANZ77411.1}; | 56 | 215 |
| tr\|A0A1B2JFK5\|A0A1B2JFK5_PICPA | Elongation factor 1-alpha {ECO:0000256\|RuleBase:RU000325}; | 49 | 205 |
| sp\|K2C1_HUMAN\| | Keratin, type II cytoskeletal 1; 67 kDa cytokeratin; Cytokeratin-1; ... | 37 | 178 |
| **PpCesA5** | **Cellulose synthase 5, glycosyltransferase family 2 [*Physcomitrella patens*]** | **37** | **155** |
| tr\|A0A1B2JH38\|A0A1B2JH38_PICPA | U3 small nucleolar RNA-associated protein 22 {ECO:0000256\|RuleBase:RU364032}; … | 34 | 132 |
| sp\|K1C10_HUMAN\| | Keratin, type I cytoskeletal 10; Cytokeratin-10; CK-10; Keratin-10; ... | 33 | 101 |
| sp\|ALBU_HUMAN\| | Albumin; Flags: Precursor; | 28 | 79 |
| tr\|A0A1B2JGF9\|A0A1B2JGF9_PICPA | BA75_03693T0 {ECO:0000313\|EMBL:ANZ77082.1}; | 20 | 71 |
| sp\|K22E_HUMAN\| | Keratin, type II cytoskeletal 2 epidermal; Cytokeratin-2e; CK-2e; … | 19 | 65 |
| sp\|K1C9_HUMAN\| | Keratin, type I cytoskeletal 9; Cytokeratin-9; CK-9; Keratin-9; … | 18 | 63 |
| tr\|A0A1B2JFJ2\|A0A1B2JFJ2_PICPA | BA75_04060T0 {ECO:0000313\|EMBL:ANZ76763.1}; | 16 | 59 |
| sp\|TRYP_PIG\| | Trypsin; EC 3.4.21.4; Flags: Precursor; | 12 | 56 |
| tr\|A0A1B2J5T9\|A0A1B2J5T9_PICPA | Plasma membrane ATPase {ECO:0000256\|RuleBase:RU362083}; EC 7.1.2.1 … | 9 | 50 |
| tr\|A0A1B2JDM7\|A0A1B2JDM7_PICPA | BA75_03179T0 {ECO:0000313\|EMBL:ANZ76146.1}; | 9 | 44 |
| tr\|A0A1B2JB83\|A0A1B2JB83_PICPA | Non-specific serine/threonine protein kinase {ECO:0000256\|SAAS:SAAS00804148}; … | 8 | 41 |
| tr\|A0A1B2JGF7\|A0A1B2JGF7_PICPA | 26S proteasome regulatory subunit RPN1 {ECO:0000256\|PIRNR:PIRNR015965}; … | 8 | 37 |
| tr\|A0A1B2JG11\|A0A1B2JG11_PICPA | BA75_03821T0 {ECO:0000313\|EMBL:ANZ76785.1}; | 7 | 36 |

| | | | |
|---|---|---|---|
| tr\|A0A1B2JEB9\|A0A1B2JEB9_PICPA | BA75_04327T0 {ECO:0000313\|EMBL:ANZ76400.1}; | 7 | 34 |
| tr\|A0A1B2JD96\|A0A1B2JD96_PICPA | BA75_02554T0 {ECO:0000313\|EMBL:ANZ75728.1}; | 7 | 34 |
| tr\|A0A1B2J6Q6\|A0A1B2J6Q6_PICPA | 5—-3— exoribonuclease 1 {ECO:0000256\|PIRNR:PIRNR006743}; EC 3.1.13.- … | 6 | 29 |
| tr\|A0A1B2JGS7\|A0A1B2JGS7_PICPA | BA75_04779T0 {ECO:0000313\|EMBL:ANZ77264.1}; | 6 | 28 |
| tr\|A0A1B2J5X6\|A0A1B2J5X6_PICPA | DNA topoisomerase I {ECO:0000256\|RuleBase:RU365101}; EC 5.6.2.1 … | 6 | 26 |
| tr\|A0A1B2JBK5\|A0A1B2JBK5_PICPA | | 5 | 25 |
| tr\|A0A1B2JCJ9\|A0A1B2JCJ9_PICPA | BA75_03101T0 {ECO:0000313\|EMBL:ANZ75528.1}; | 5 | 22 |
| tr\|A0A1B2JEC7\|A0A1B2JEC7_PICPA | S-adenosylmethionine synthase {ECO:0000256\|RuleBase:RU000541}; … | 5 | 22 |
| tr\|A0A1B2JFH4\|A0A1B2JFH4_PICPA | BA75_03691T0 {ECO:0000313\|EMBL:ANZ76819.1}; | 5 | 21 |

**Supplementary Table S5.** List of *Pichia pastoris* proteins in the excised single band (~110 kDa) corresponding to PttCesA8 as identified by mass spectrometry.

| **Protein** | **Description** | **Unique** | **Total** |
|---|---|---|---|
| tr\|A0A1B2JF91\|A0A1B2JF91_PICPA | BA75_03713T0 {ECO:0000313\|EMBL:ANZ76707.1}; | 474 | 947 |
| **PttCesA8** | **Cellulose synthase 8, glycosyltransferase family 2 [*Populous tremuloides*]** | **54** | **225** |
| sp\|K1C10_HUMAN\| | Keratin, type I cytoskeletal 10; Cytokeratin-10; CK-10; Keratin-10; ... | 51 | 212 |
| sp\|K2C1_HUMAN\| | Keratin, type II cytoskeletal 1; 67 kDa cytokeratin; Cytokeratin-1; ... | 43 | 188 |
| sp\|K22E_HUMAN\| | Keratin, type II cytoskeletal 2 epidermal; Cytokeratin-2e; CK-2e; ... | 38 | 155 |
| tr\|A0A1B2JFJ2\|A0A1B2JFJ2_PICPA | BA75_04060T0 {ECO:0000313\|EMBL:ANZ76763.1}; | 73 | 146 |
| tr\|A0A1B2JB82\|A0A1B2JB82_PICPA | BA75_02757T0 {ECO:0000313\|EMBL:ANZ75231.1}; | 53 | 107 |
| sp\|TRYP_PIG\| | Trypsin; EC 3.4.21.4; Flags: Precursor; | 15 | 85 |
| sp\|K1C9_HUMAN\| | Keratin, type I cytoskeletal 9; Cytokeratin-9; CK-9; Keratin-9; ... | 19 | 81 |
| tr\|A0A1B2JGC1\|A0A1B2JGC1_PICPA | BA75_04232T0 {ECO:0000313\|EMBL:ANZ77043.1}; | 38 | 75 |
| sp\|P02769\|ALBU_BOVIN | Serum albumin; BSA; Bos d 6; Flags: Precursor; | 36 | 70 |
| tr\|A0A1B2JHN5\|A0A1B2JHN5_PICPA | BA75_05137T0 {ECO:0000313\|EMBL:ANZ77411.1}; | 32 | 64 |
| tr\|A0A1B2JHU5\|A0A1B2JHU5_PICPA | Protein kinase C {ECO:0000256\|SAAS:SAAS01199224}; EC 2.7.11.13 ... | 31 | 61 |
| tr\|A0A1B2JG55\|A0A1B2JG55_PICPA | 5—-3— exoribonuclease {ECO:0000256\|PIRNR:PIRNR037239}; EC 3.1.13.- ... | 30 | 60 |
| tr\|A0A1B2JEB9\|A0A1B2JEB9_PICPA | BA75_04327T0 {ECO:0000313\|EMBL:ANZ76400.1}; | 27 | 54 |
| tr\|A0A1B2JG25\|A0A1B2JG25_PICPA | BA75_04202T0 {ECO:0000313\|EMBL:ANZ77009.1}; | 26 | 51 |
| tr\|A0A1B2J5T9\|A0A1B2J5T9_PICPA | Plasma membrane ATPase {ECO:0000256\|RuleBase:RU362083}; EC 7.1.2.1 ... | 24 | 47 |
| tr\|A0A1B2JGF9\|A0A1B2JGF9_PICPA | BA75_03693T0 {ECO:0000313\|EMBL:ANZ77082.1}; | 22 | 46 |
| tr\|A0A1B2JH86\|A0A1B2JH86_PICPA | BA75_04399T0 {ECO:0000313\|EMBL:ANZ77424.1}; | 22 | 44 |

| | | | |
|---|---|---|---|
| tr\|A0A1B2JH38\|A0A1B2JH38_PICPA | U3 small nucleolar RNA-associated protein 22 {ECO:0000256\|RuleBase:RU364032}; ... | 24 | 44 |
| tr\|A0A1B2JC36\|A0A1B2JC36_PICPA | NAD-specific glutamate dehydrogenase {ECO:0000256\|PIRNR:PIRNR000184}; ... | 20 | 39 |
| tr\|A0A1B2JEB1\|A0A1B2JEB1_PICPA | BA75_03282T0 {ECO:0000313\|EMBL:ANZ76185.1}; | 19 | 38 |
| tr\|A0A1B2JB87\|A0A1B2JB87_PICPA | BA75_02308T0 {ECO:0000313\|EMBL:ANZ75249.1}; | 18 | 36 |
| tr\|A0A1B2J9Q0\|A0A1B2J9Q0_PICPA | BA75_00239T0 {ECO:0000313\|EMBL:ANZ74741.1}; | 18 | 35 |
| tr\|A0A1B2JFK5\|A0A1B2JFK5_PICPA | Elongation factor 1-alpha {ECO:0000256\|RuleBase:RU000325}; | 16 | 32 |
| sp\|RS27A_HUMAN\| | Ubiquitin-40S ribosomal protein S27a; Ubiquitin carboxyl extension ... | 15 | 31 |
| tr\|A0A1B2JG11\|A0A1B2JG11_PICPA | BA75_03821T0 {ECO:0000313\|EMBL:ANZ76785.1}; | 14 | 29 |
| sp\|ALBU_HUMAN\| | Albumin; Flags: Precursor; | 7 | 14 |
| tr\|A0A1B2JEY5\|A0A1B2JEY5_PICPA | BA75_03358T0 {ECO:0000313\|EMBL:ANZ76553.1}; | 12 | 23 |
| tr\|A0A1B2JDM7\|A0A1B2JDM7_PICPA | BA75_03179T0 {ECO:0000313\|EMBL:ANZ76146.1}; | 11 | 22 |

**Supplementary Text S1.** Protein sequences of Cellulose Synthases used in this study. All the tags in the proteins are underlined. PpCesA5 has a C-terminal 12x-HIS tag, and PttCesA8 has an N-terminal FLAG tag and a C-terminal 12x-HIS tag.

>>PpCesA5

MEANAGLIAGSHNRNELVVLRPDHEGPKPLSQVNSQFCQICGDDVGVTVDGELFVACF
ECGFPVCRPCFEYERKEGNQSCPQCKSRYNRQKGSPRVPGDEEEDDTDDLENEFALEMG
QLDEQNVTDAMLHGHMSYGGNYDHNLPNLHQTPQFPLLTDGKMGDLDDDSHAIVLPP
PMNGGKRVHPLPYIESNLPVQARPMDPTKDLAAYGYGSVAWKDRVESWKMRQEKMM
TEGSHHHKGGDMDGDNGPDLPIMDEARQPLSRKVPISSARINPYRMLIVIRLVVLAFFFR
YRILNPVEGAYGMWLTSVICEIWFAISWILDQFPKWLPINRETYLDRLSLRYEKEGEPSQ
LEHVDIFVSTVDPMKEPPLVTANTILSILAVDYPVDKVSCYLSDDGAAMLTFECISETSEF
ARKWVPFCKKFSIEPRAPEMYFAQKIDYLKDKVQPTFVKERRAMKREYEEFKVRVNAL
VAKAQKVPEEGWTMQDGTPWPGNNSRDHPGMIQVFLGHSGGHDTDGNELPRLVYVSR
EKRPGFNHHKKAGAMNALVRVSAVLTNAPYFLNLDCDHYINNSKALREAMCFFMDPS
VGKKVCYVQFPQRFDGIDRNDRYANHNTVFFDINLKGLDGIQGPVYVGTGTVFNRKAL
YGYEPVLKEKESKGTGCGAACSTLCCGKRKKDKKKNKKSKFSRKKTAPTRSDSNIPIFS
LEEIEEGDEEKSSLVNTINYEKRFGQSPVFVASTLLEHGGVHHSASPGSLLKEAIHVISCG
YEDKTDWGKEIGWIYGSVTEDILTGFKMHCRGWRSIYCMPTRPAFKGSAPINLSDRLNQ
VLRWALGSVEISLSRHCPLWYGYGGRLKCLERLAYINTTIYPLTSLPLVAYCVLPAVCLL
TGNFIIPTISNLDSLYFISLFLSIFVTGILEMRWSGVGIDEWWRNEQFWVIGGVSAHLFALF
QGLLKVFAGVDTNFTVTSKQADDEDFGELYMLKWTSLLIPPTTILILNLVGVVAGISDAI
NNGYQSWGPLFGKLFFAFWVIVHLYPFLKGLMGRQNRTPTIVIVWSILLASIFSLLWVRI
NPFLSRSNGPNLVECGLSCHHHHHHHHHHHH

>>PttCesA8

MDYKDDDDKYPYDVPDYAMMESGAPICHTCGEQVGHDANGDLFVACHECNYHICKS
CFEYEIKEGRKVCLRCGSPYDENLLDDVEKKGSGNQSTMASHLNNSQDVGIHARHISSV
STVDSEMNDEYGNPIWKNRVESWKDKRNKKKKSNTKPETEPAQVPPEQQMENKPSAE
ASEPLSIVYPIPRNKLTPYRAVIIMRLIILGLFFHYRITNPVDSAFGLWLTSVICEIWFAFSW
VLDQFPKWKPVNRETFIERLSARYEREGEPSQLAAVDFFVSTVDPLKEPPLITANTVLSIL
AVDYPVDKVSCYVSDDGAAMLTFESLVETAEFARKWVPFCKKFSIEPRAPEFYFSQKID
YLKDKVQPSFVKERRAMKRDYEEYKVRVNALVAKAQKTPDEGWTMQDGTPWPGNNT
RDHPGMIQVFLGNTGARDIEGNELPRLVYVSREKRPGYQHHKKAGAENALVRVSAVLT
NAPYILNLDCDHYVNNSKAVREAMCILMDPQVGRDVCYVQFPQRFDGIDRSDRYANRN
IVFFDVNMKGLDGIQGPMYVGTGCVFNRQALYGYGPPSMPRLRKGKESSSCFSCCCPTK
KKPAQDPAEVYRDAKREDLNAAIFNLTEIDNYDDYERSMLISQLSFEKTFGLSPVFIESTL
MENGGVPESANSSTLIKEAIHVIGCGFEEKTEWGKEIGWIYGSVTEDILSGFKMHCRGWR
SIYCMPVRPAFKGSAPINLSDRLHQVLRWALGSVEIFFSRHCPFWYGYGGRLKWLQRL
AYINTIVYPFTSLPLIAYCTIPAVCLLTGKFIIPTLSNLASMLFLGLFISIIVTAVLELRWSGV
SIEDLWRNEQFWVIGGVSAHLFAVFQGFLKMLAGIDTNFTVTAKAADDTEFGELYMVK
WTTLLIPPTTLLIINIVGVVAGFSDALNKGYEAWGPLFGKVFFAFWVILHLYPFLKGLMG
RQNRTPTIVVLWSVLLTSVFSLVWVKINPFVNKVDNTLAGETCISIDCHHHHHHHHHHH
H
